# Inferring Fitness Landscapes and Selection on Phenotypic States from Single-Cell Genealogical Data

**DOI:** 10.1101/069260

**Authors:** Takashi Nozoe, Edo Kussell, Yuichi Wakamoto

## Abstract

Recent advances in single-cell time-lapse microscopy have revealed non-genetic heterogeneity and temporal fluctuations of cellular phenotypes. While different phenotypic traits such as abundance of growth-related proteins in single cells may have differential effects on the reproductive success of cells, rigorous experimental quantification of this process has remained elusive due to the complexity of single cell physiology within the context of a proliferating population. We introduce and apply a practical empirical method to quantify the fitness landscapes of arbitrary phenotypic traits, using genealogical data in the form of population lineage trees which can include phenotypic data of various kinds. Our inference methodology for fitness landscapes determines how reproductivity is correlated to cellular phenotypes, and provides a natural generalization of bulk growth rate measures for single-cell histories. Using this technique, we quantify the strength of selection acting on different cellular phenotypic traits within populations, which allows us to determine whether a change in population growth is caused by individual cells’ response, selection within a population, or by a mixture of these two processes. By applying these methods to single-cell time-lapse data of growing bacterial populations that express a resistance-conferring protein under antibiotic stress, we show how the distributions, fitness landscapes, and selection strength of single-cell phenotypes are affected by the drug. Our work provides a unified and practical framework for quantitative measurements of fitness landscapes and selection strength for any statistical quantities definable on lineages, and thus elucidates the adaptive significance of phenotypic states in time series data. The method is applicable in diverse fields, from single cell biology to stem cell differentiation and viral evolution.

## Introduction

Selection is a process in which the interaction of organisms with their environment determines which types of individuals thrive and proliferate more than others. Genetic information encoded in the genome is a primary determinant of reproductivity, but epigenetic and fluctuating phenotypic traits can also strongly influence selection [1–4]. Recent single-cell measurements revealed the existence of phenotypic heterogeneity within clonal populations, including cases in which heterogeneity has been shown to have a clear functional role [5, 6] such as bacterial persistence [7–9], infection [10], and competence and sporulation [11]. Quantifying reproductivity of phenotypic traits and revealing how strongly selection acts within a clonal population are thus of crucial importance for understanding the biological significance of phenotypic heterogeneity.

To experimentally evaluate reproductivity of a unicellular organism, one usually measures bulk growth rate (Malthusian parameter [12]) of a cellular population in batch or uses a competition assay between a genotype of interest and a reference genotype [13]. These methods are only valid when the time-scale of genotypic changes is sufficiently long compared with that of the measurements. However, the time-scale of phenotypic changes is often comparable to cellular generation time ([14] and Fig S1), and only in certain cases is it orders of magnitude longer, e.g. when a phenotypic state is stabilized by specific epigenetic and/or positive-feedback regulations. As a result, bulk population growth rates of sub-populations fractionated based on initial phenotypic traits, e.g. by fluorescence-activated cell sorting, do not necessarily represent reproductivity of initial phenotypic traits because phenotypic traits are diversified rapidly by complex dynamical processes that occur during measurements. An alternative approach is necessary to measure reproductivity for heterogeneous and fluctuating cellular phenotypes.

Using time-lapse microscopy and fluorescent reporters, it has become possible to follow full individual cell histories recording all division events and instantaneous expression levels of reporters within cellular populations [8, 15–20]. Several theoretical studies have demonstrated the utility of history-based analysis of growing populations, regarding individual histories rather than single cells as the basic replicating entity [21–23]. For example, Leibler and Kussell introduced a time-integrated instantaneous reproduction rate, termed *historical fitness* [21], and defined a measure of selection using the response of mean historical fitness over all histories within a population. However, empirically determining the instantaneous reproduction rate of an individual cell can be difficult in general, e.g. due to the fact that cell size, age, elongation rate, and division timing are a subset of possible observables all of which contribute to reproduction. Evaluating the fitness value of a certain phenotypic trait such as expression level of a specific gene results in additional complications.

To address these difficulties, we introduce an empirically measurable quantity associated with phenotypic states, which we call the phenotypic *fitness landscape*. This quantity, which reports how cellular reproductivity is correlated with phenotypic states, extends the definition of historical fitness so that it becomes meaningful in a general setting without requiring any assumptions. Our approach allows one to assign a fitness value to any statistical quantities observed over cellular lineages, and to evaluate the selection strength acting on different phenotypic states. To formulate our framework, we leverage a fundamental property of selection processes: the *retrospective probability* of observing a certain phenotypic trait value by moving backward in time from the present to the ancestral parts of a lineage is different from its counterpart, the *chronological probability* to observe the trait value moving forward in time along a lineage as individuals grow and divide. We show that these two probabilities can be evaluated directly using single-cell lineage tree data, leading to natural definitions of the fitness landscape and selection strength. We apply this framework to analyze proliferation processes of simulated and experimental cellular populations, demonstrating the utility of our measures to reveal phenotype dependent fitness and its response to environmental change.

## Results

### Lineage trees and transitions of phenotypes along cell lineages

We first present an overview of the type of biological data that is examined in this study. Figure 1A shows an example of time-lapse images of *E. coli* growing on an agarose pad. Analyzing the images provides information on the proliferation dynamics and phenotype transitions (Fig 1B–1D). For example, one can reveal the lineage tree structure, which shows genealogical relationships among the individual cells that originated from an ancestor cell (Fig 1B). One can also extract the information on the transitions of cell size along cell lineages (Fig 1C) and of other cellular phenotypes such as intracellular concentration of a particular protein if it can be probed with an appropriate fluorescence reporter (Fig 1D). Here we regard any measurable quantity or set of quantities observed along cell lineages as a quantifiable phenotype in the broadest sense. We note that this type of information is now available for many biological processes including embryogenesis and stem cell differentiation [19,20,24–27]. As shown in Fig 1B, division intervals (i.e. cell cycle durations) of individual cells are usually heterogeneous, yielding variability in the number of divisions observed along different cell lineages. Moreover, cellular phenotypes also fluctuate along cell lineages (Fig 1C and 1D). To understand the role of such phenotypic heterogeneity for population growth, one must know whether the difference of the phenotypic states are correlated with the growth rate heterogeneity within a population. Below we present the theoretical results and the analytical procedure that allow us to reveal the quantitative relations between phenotypes and growth.

**Fig 1.**
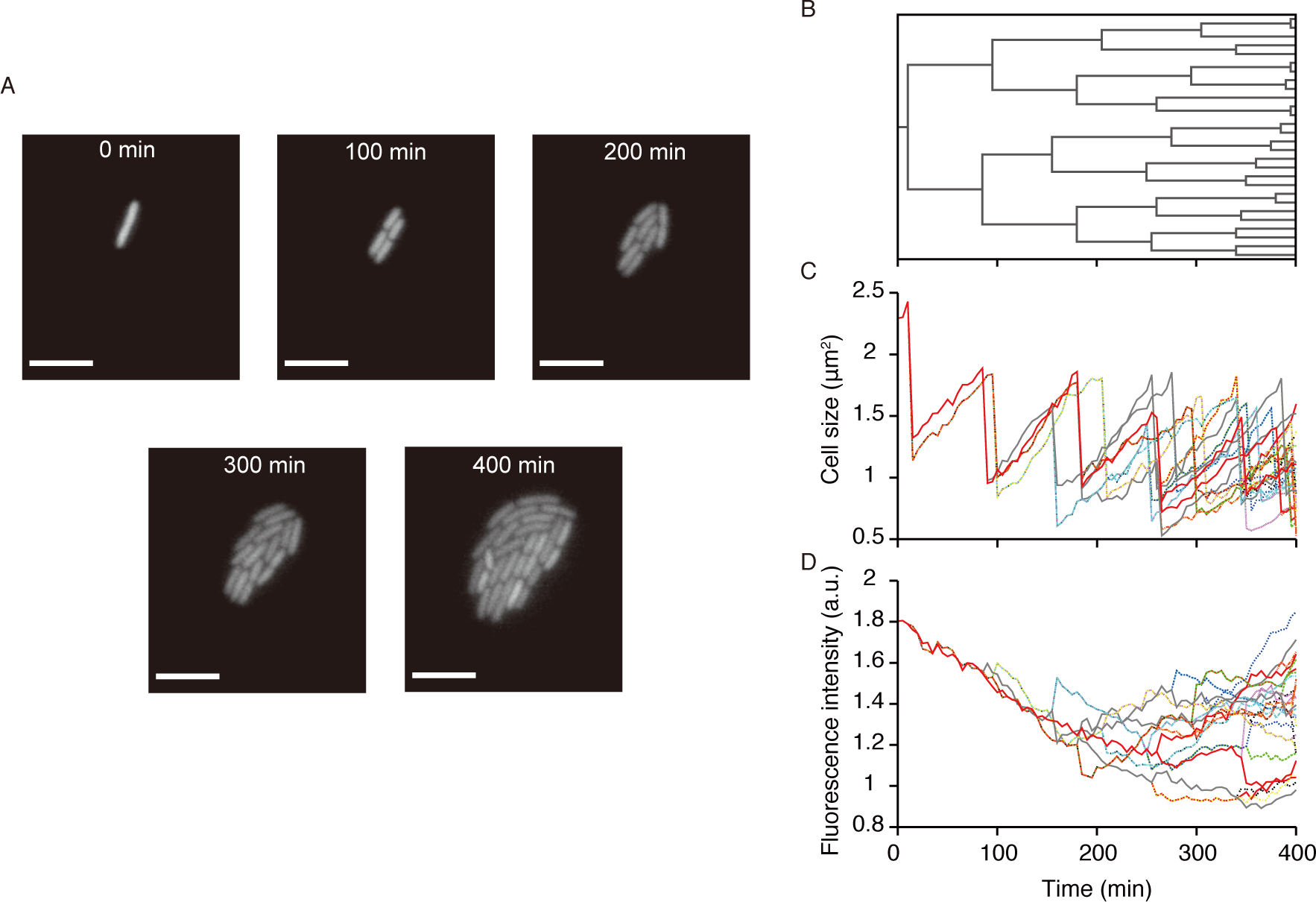
An example of cell lineage data obtained by single-cell time-lapse measurement. **A** Time-lapse images of a growing microcolony of *E. coli* F3/pTN001. This strain expresses a fluorescent protein, Venus-YFP, from a low copy plasmid (see Materials and Methods). One can obtain the time-lapse images for many microcolonies (ca. 100) starting from different ancestral cells in a single experiment. Scale bars, 5 µm. **B** Cell lineage tree for the microcolony in (A), which shows the genealogical relationship of individual cells. **C** Transitions of cell sizes. different lines correspond to different single-cell lineages on the tree (B). Due to the fluctuation of growth parameters such as elongation rate and division interval, the transition patterns are variable among the cell lineages. **D** Transitions of mean fluorescence intensity along single-cell lineages on the tree (B). Again, the transition patterns are variable among the cell lineages.

### Retrospective and chronological probabilities for single cell lineages

We consider a binary division process as depicted in Fig 2, where *t*_0_, *t*_1_ are the start and end times of a lineage tree, and we define *τ* = *t*_1_ − *t*_0_ as the duration of observation. To illustrate our view of lineage statistics, we first consider a single fixed lineage tree denoted by 𝒯 derived from a single ancestor cell (Fig 2A). Let *N* (*t*, T) be the number of cells in the tree 𝒯 at time *t* and we label and distinguish each lineage by *i* = 1, 2,…, *N* (*t*_1_, 𝒯). For the tree 𝒯 in Fig 2A, *N* (*t*_1_, 𝒯) = 13. We consider two different ways of randomly sampling single-cell lineages on the tree. We could sample each lineage with equal weight, where the probability of choosing lineage *i* is 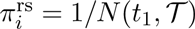, which we call the *retrospective probability* because it corresponds to the probability that the past history of the last cell on lineage *i* is chosen. For the tree 𝒯, 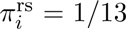 for all *i*. Alternatively, letting *D_i_* be the number of cell divisions on lineage *i*, we could sample lineage *i* with probability 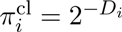, which we call the *chronological probability* because it is the probability that lineage *i* is chosen by descending the tree from the ancestor cell at *t*_0_ randomly at each branch point with equal probability 1/2. For examples, 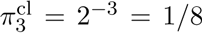 and 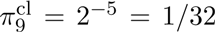 for the tree 𝒯 (Fig 2A). The probability distribution 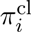 is determined solely by the number of divisions on lineage *i*, being unaffected by the reproductive performance of the other lineages. In contrast, 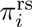 strongly depends on the reproductive performance of the other lineages, which enters into the total number of cell lineages *N* (*t*_1_, 𝒯). Generally, the ratio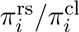 is positively correlated with the relative reproductive performance of lineage *i*. In fact, 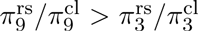 for 𝒯. In addition, we note that the inconsistencies between 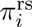 and 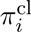 reflect the variability in the numbers of divisions among the cell lineages. It can be confirmed that 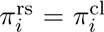 for all *i* if the same number of divisions occur for all the lineages on a tree (Fig 2B).

**Fig 2.**
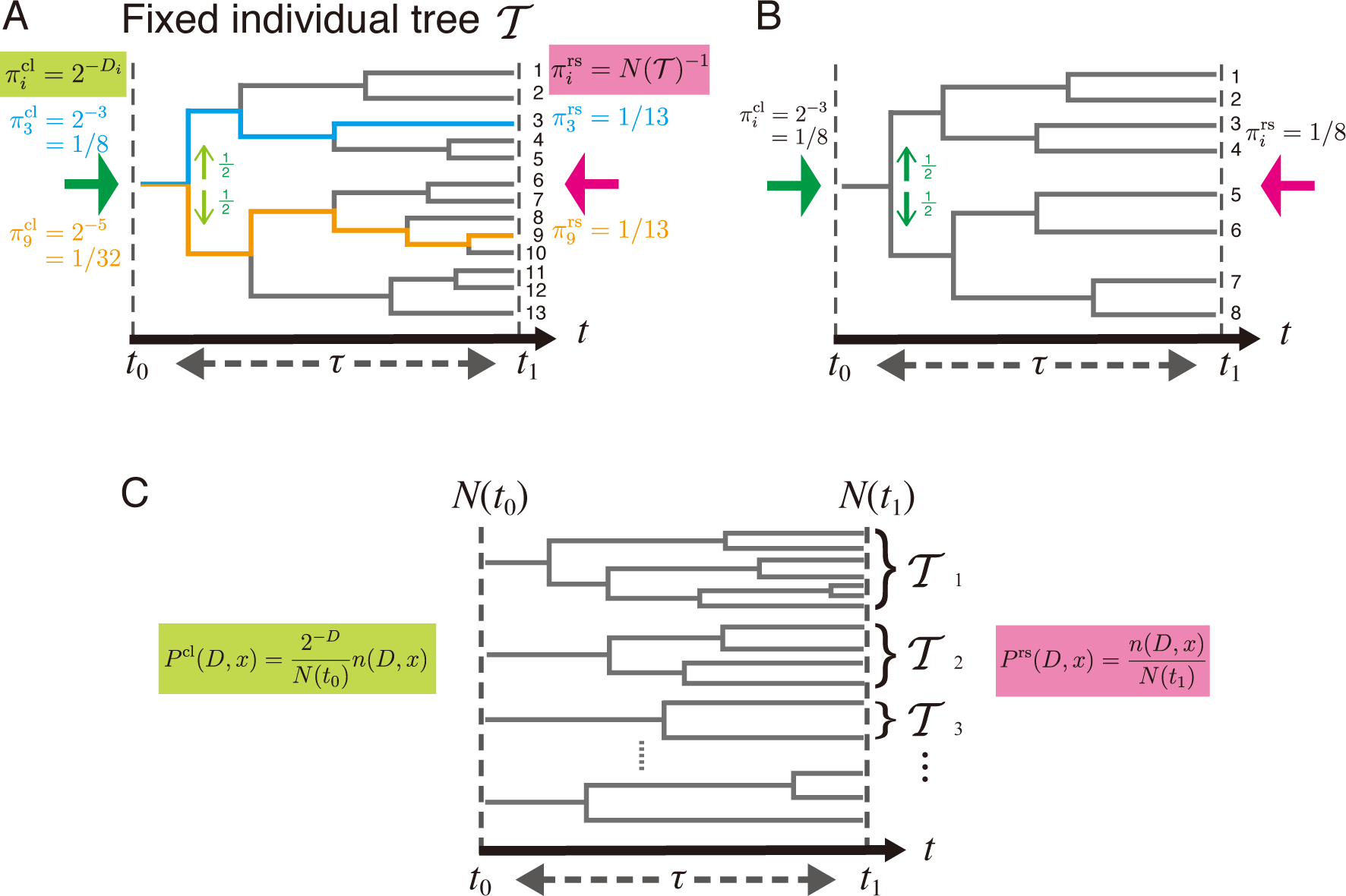
Chronological and retrospective probabilities of single-cell lineages. A. Chronological and retrospective probabilities on a fixed tree. Here we consider a representative fixed lineage tree 𝒯 spanning from time *t*_0_ to *t*_1_ = *t*_0_ + *τ*. The number of cells in this tree at *t*_1_ is *N* (*t*_1_, 𝒯) = 13 cells, and each of these cells distinguishes a unique lineage (e.g. the cyan and orange lines in the tree). 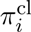 is the probability that a cell lineage *i* (*i* = 1, 2, · · ·, *N* (*t*_1_, 𝒯)) is chosen by descending the tree from *t*_0_ to *t*_1_ (green arrow). At every division point, we randomly select one daughter cell’s lineage with the probability of 1/2 (light green arrows). The probability that we choose lineage *i* in this manner is 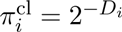, where *D_i_* is the number of cell divisions on lineage *i*. 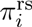 is the probability of choosing cell lineage *i* among *N* (*t*_1_, T) lineages with equal weight (pink arrow). Thus, 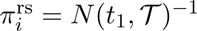. We call 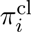 and 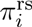 the chronological probability and retrospective probability, respectively, based on the time directions of the green and pink arrows. The chronological and retrospective probabilities for the cell lineages 3 and 9 are shown in cyan and red texts, respectively. **B.** A tree on which all the cell lineages have the same number of cell divisions. In this case, the chronological and the retrospective probabilities are equal for all the lineages. **C.** General case with a large collection of lineage trees. T_*i*_ denotes a tree each descended from a different ancestor cell at time *t*_0_. The definitions of the chronological and retrospective joint probabilities of division count *D* and lineage phenotype *x* are shown in green and pink, respectively. *n*(*D, x*) denotes the total number of cell lineages with *D* and *x*, i.e. 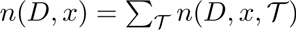.

### Lineage fitness and fitness landscape on a phenotypic trait

The consideration above indicates that the cell lineages with more divisions are over-represented in the retrospective probability relative to its chronological probability. This idea can be further extended to the general situation where a large number of lineage trees are contained in the population. We now consider the set of lineages within a large collection of independent trees initiated from a large number of progenitor cells *N* (*t*_0_) » 1 (Fig 2C). For each lineage, we record a phenotypic trait *x* and the number of divisions *D*, where *x* can be any random variable representing a phenotypic trait of a single cell lineage, e.g. a time-averaged gene expression level, average cell length, number of divisions *D*, or any variety of other possibilities. We consider the joint statistics of *D* and *x* across all possible trees, letting *n*(*D, x*, 𝒯) denote the number of lineages with values *D* and *x* within tree 𝒯, and we denote the sum of this quantity over trees as *n*(*D, x*). The total number of lineages observed across all trees, *N* (*t*_1_), is given by summing *n*(*D, x*) over *D* and *x* (see Supplemental Information). In analogy with the single tree quantities, we define the retrospective probability of choosing a lineage with *D* and *x* as

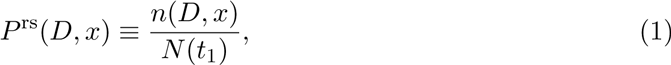

and the chronological probability as

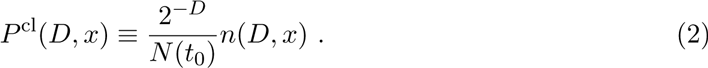

Defining Λ to be the population growth rate,

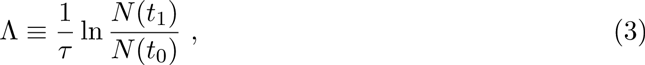

we obtain using equations 1 and 2 the relation

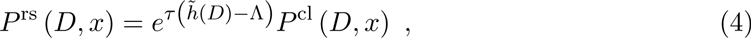

where *h*˜(*D*) ≡ *τ*^*−*1^*D* ln 2. We see from Eq. 4 that *h*˜(*D*) is the natural measure of fitness for a lineage, since lineages for which this quantity is greater than Λ will be exponentially over-represented in retrospective probability relative to chronological probability. We call *h*˜(*D*) the *lineage fitness*.

We now measure how quickly the number of lineages with a given phenotype *x* grow between times *t*_0_ and *t*_1_ according to their chronological and retrospective probabilities. We denote by *P*^rs^(*x*) ≡ ∑_*D*_ *P*^rs^(*D, x*) and *P*^cl^(*x*) ≡ ∑_*D*_ *P*^cl^(*D, x*) the retrospective and chronological marginal probability distributions of *x*. We define the phenotypic *fitness landscape h*(*x*) as

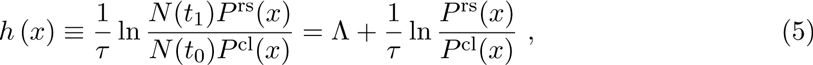

noting that *N* (*t*_0_)*P*^cl^(*x*) and *N* (*t*_1_)*P*^rs^(*x*) are the effective numbers of cell lineages with a phenotypic trait *x* from the chronological and retrospective perspectives, respectively. We can rewrite Eq. 5 as

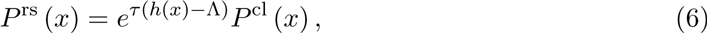

which shows that if *h*(*x*) is greater than Λ the phenotypic state *x* will be exponentially over-represented in retrospective relative to chronological probability. Thus, *h*(*x*) provides a natural extension of fitness for lineage-based phenotypic traits. We point out that both *P*^rs^(*D, x*) and *P*^cl^(*D, x*) (hence, *P*^rs^(*x*) and *P*^cl^(*x*) as well) are obtainable directly from the set of lineage trees (Fig 2C). Thus, *h*(*x*) can also be determined directly from the lineage tree data using Eq. 5.

In Fig 3, we schematically show how the fitness landscapes look depending on the deviation between chronological and retrospective probability distributions. When the deviation is small, the fitness landscape *h*(*x*) is flat over the phenotypic space *x* (Fig 3A); when the deviation is large, *h*(*x*) changes greatly depending on *x* (Fig 3B). In the next section, we quantify the amount of heterogeneity in *h*(*x*) using the selection strength, *S*[*x*], defined below.

**Fig 3.**
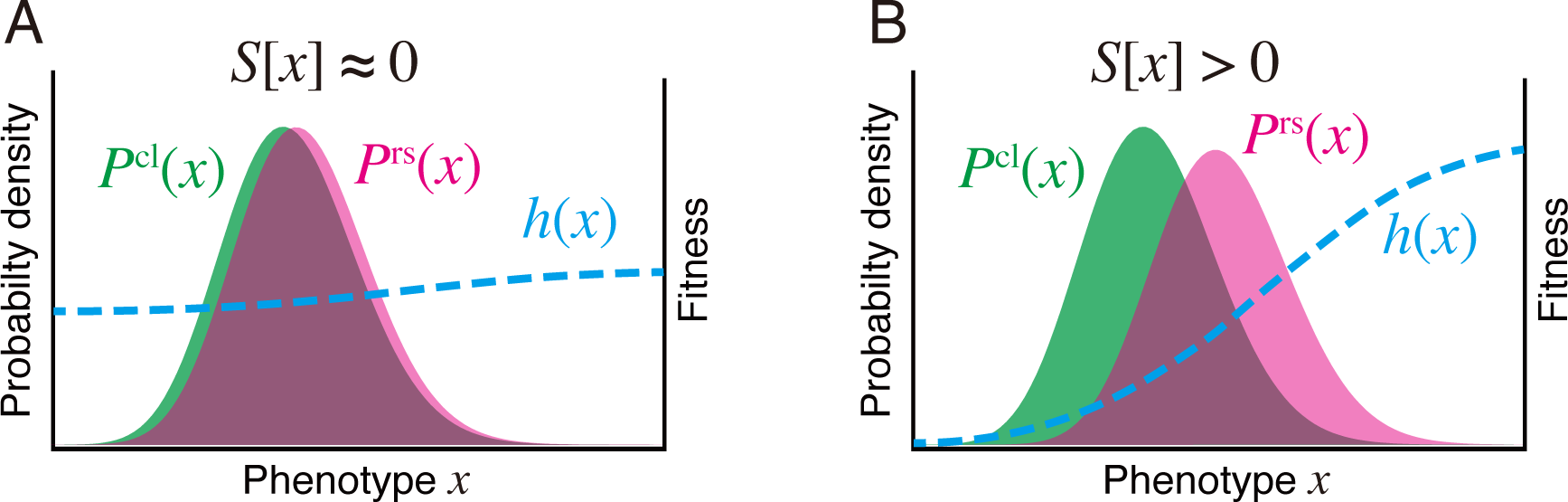
Schematic pictures that show the relations among chronological and retrospective phenotype distributions, fitness landscape, and selection strength. **A.** Weak selection. When the chronological (green) and retrospective (pink) probability distributions are similar, the fitness landscape *h*(*x*) largely does not change over the range of phenotypic values *x*, and the selection strength *S*[*x*] is approximately 0. **B.** Strong selection. When the retrospective probability distribution significantly deviates from the chronological distribution, the fitness landscape *h*(*x*) changes greatly over the range of phenotypic values *x*. In this case, the selection strength *S*[*x*] is greater than zero.

### Measuring the strength of phenotypic selection

Specific states of the phenotypic trait *x* can be selected if *x* and *D* are correlated. In general, the strength of this correlation could differ significantly among different phenotypes. In the conventional framework of natural selection known as Fisher’s fundamental theorem, selection strength is measured by the gain of mean fitness due to the change of probability distribution of a phenotype [12]. Inspired by this idea, we define the strength of selection acting on a phenotypic trait *x* as

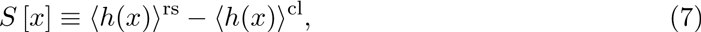

where 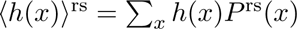 and 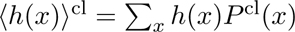 are the mean fitness in retrospective and chronological perspectives, respectively.

This simple measure of selection strength has rich underpinnings. First, *S*[*x*] is also a measure of fitness variation on the landscape *h*(*x*) because

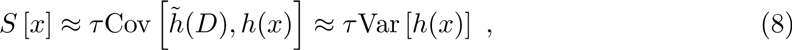

where the variance and covariance can equivalently be taken over either chronological or retrospective distributions, and the approximation is accurate to the order of second cumulants of *h*˜(*D*) and *h*(*x*) (see *Supporting Information*). Therefore, *S*[*x*] ≈ 0 if *h*(*x*) is uniform over the range of observed value of *x*, but *>* 0 if *h*(*x*) changes significantly with *x* (Fig 3). Secondly, *S*[*x*] also represents the statistical deviation between the probability distributions *P*^cl^(*x*) and *P*^rs^(*x*) because

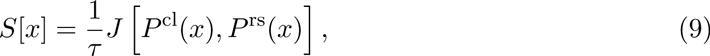

where 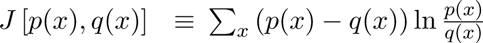 is the Jeffereys divergence [28–30], a non-negative quantity that measures the dissimilarity between two probability distributions (see *Supporting Information*). From the properties of Jeffreys divergence, we can prove that

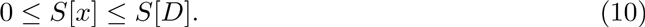

The strength of selection acting on any phenotypic state is therefore bounded by the strength of selection acting on *D*, i.e. the maximal possible value. As described in *Supporting Information, S*[*x*] can be interpreted as an amount of information representing to what extent variation of *D* can be explained by phenotype *x*. Therefore, when *S*[*x*] is large, phenotype *x* is strongly correlated with lineage fitness. In fact, we prove that the *relative selection strength*, defined by

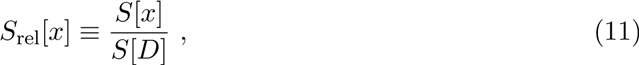

is approximately equal to the squared correlation coefficient between *h*˜(*D*) and *h*(*x*) to the order of second cumulants (Eq. S1.55).

### Decomposition of fitness response to environmental change

We now introduce an explicit dependence of all quantities on an environment variable ε, and using this notation Eq. 7 becomes

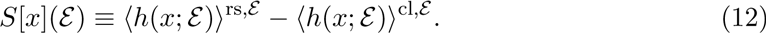

Let us denote the changes of mean fitness and selection strength due to an environmental shift from *ε*_1_ to *ε*_2_ as 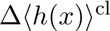, 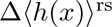 and Δ*S*[*x*]. Then

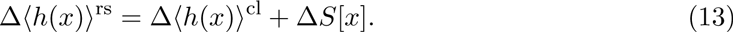

Δ〈*h*(*x*)〉^rs^ represents the response of mean fitness in retrospective histories due to the change of the environments. Eq. 13 indicates that this term can be decomposed into two terms: Δ〈*h*(*x*)〉^cl^, which represents the intrinsic response to the environmental change; and Δ*S*[*x*], the change of selection strength. Thus, this framework allows us to distinguish and evaluate the contributions of individual response and selection to the total change of retrospective mean fitness.

In *Supporting Information*, we apply the above framework to several analytically tractable models, and directly calculate the fitness landscape and selection strength in each model. We also provide examples of the fitness decomposition in *Supporting Information*.

### Simulation

To demonstrate the utility of our lineage-based analysis, we first applied it to simulation data of a cell proliferation model. In this model, we consider a population in which cells divide according to division probability *f* (*y*^*t*^)Δ*t*, where Δ*t* is time increment, and *y_t_* is a variable that represents an instantaneous state of a certain phenotype at time *t* (Fig 4A). For example, *y_t_* could be the intracellular concentration of a protein of interest. In the simulation, we assume that ln *y_t_* follows the Ornstein-Uhlenbeck process so that the stationary distribution of *y_t_* in chronological cell histories follows the log-normal distribution with mean 1 and standard deviation 0.3 (Fig 4B). We set *f* (*y*) to be a Hill function, 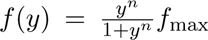, where *n* is the Hill coefficient, and *f*_max_ is the maximum division rate (Fig 4B). We fixed *f*_max_ = 1.2 h^*−*1^ and ran the simulation under different values of *n*. The initial state of a cell lineage at *t*_0_ was randomly sampled from the stationary log-normal distribution. In each condition, we repeated the simulation 100 times, i.e. *N* (*t*_0_) = 100, which is a realistic sample size of single-cell time-lapse experiments. To calculate the fitness landscape and selection strength, we used the lineage tree data between *t*_0_ = 0 min and *t*_1_ = 250 min (thus *τ* = 250 min). Additional details of the simulation are described in Materials and Methods.

**Fig 4.**
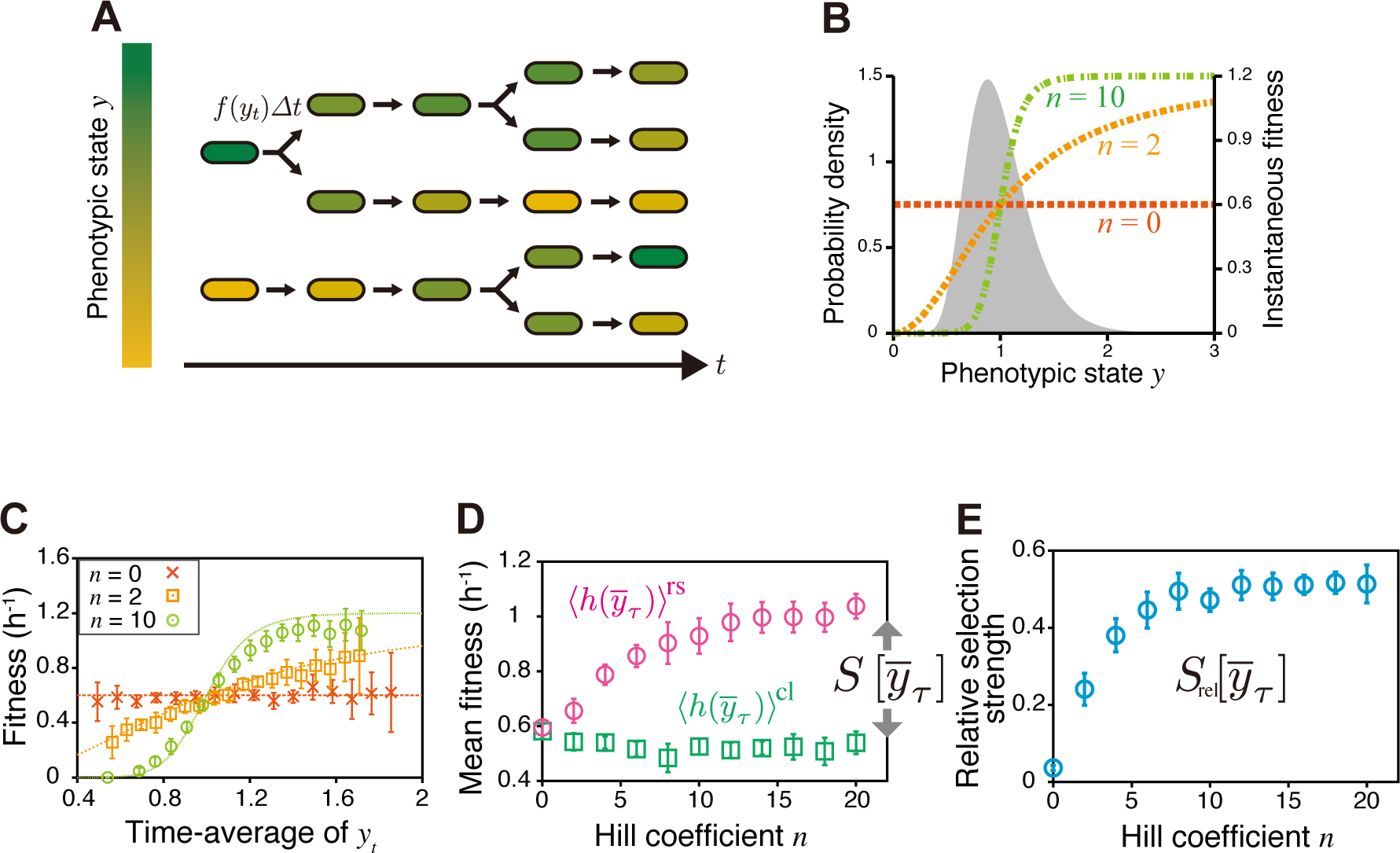
Quantifying fitness landscape and selection strength for the simulation data of clonal cell proliferation. **A.** Cell proliferation model with phenotype fluctuations. Individual cells divide according to the instantaneous fitness *f* (*y_t_*), which depends on the current phenotypic state *y_t_* (different colors indicate different phenotypic states). *y_t_* fluctuates in time, causing fitness fluctuations on single-cell lineages. **B.** Stationary probability of the phenotypic state without selection (filled curve, log-normal distribution with mean 1 and standard deviation 0.3) and instantaneous fitness depending on phenotypic states (Hill function, colored dashed lines with different Hill coeffcients, *n* = 0, 2 and 10). **C.** Fitness landscapes. We produced the datasets of clonal cell proliferation by simulation, in which we assumed that cells stochastically change phenotypic state *y* and divide in a phenotype-dependent manner with the division rate 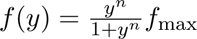. We calculated fitness landscapes 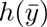 from the simulation data for the conditions of *n* = 0, 2, and 10. In all the conditions, 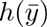 (points) recovered the assigned phenotype-dependent division rate *f* (*y*) (broken curves) with good precision. The points and the error bars represent means and standard deviations of results from 10 independent simulations (same in B and C). **D.** Dependence of mean fitness 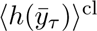 and 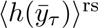 on Hill coeffcient. Strengthening the phenotype dependence of fitness by increasing the Hill coefficient of *f* (*y*) caused 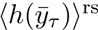 (magenta circles) to be greater than 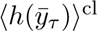 (green squares). In our definition, selection strength for phenotype 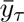 is given by 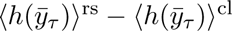 (Eq. 7), thus the deviation directly indicates the existence of selection acting on phenotype 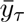. **E.** Dependence of relative selection strength 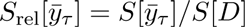 on Hill coefficient.

We tested our methodology using the time-averaged expression level as a simple phenotypic trait *x* of cell lineages, i.e.

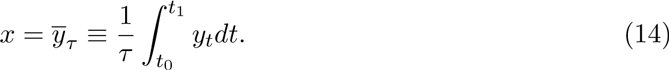

We found that the fitness landscape 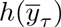 calculated from the simulated lineage trees and the time-series of *y_t_* recovers *f* (*y*) accurately despite the non-linearity of this function (Fig 4C and Fig S2A,B). The chronological mean fitness 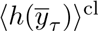 is unchanged by the change of *n*, but the retrospective mean fitness 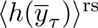 increases significantly with *n* (Fig 4D). As a result, selection strength 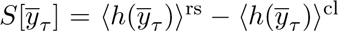 as well as relative selection strength 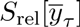 increase with *n* as expected from the fact that larger *n* introduces greater fitness variation (Fig 4D,E). Reducing the autocorrelation time of *y_t_* decreases selection strength (Fig S2C), since faster fluctuations of the phenotype decrease the variation of the time average, 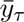. In this case, 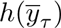 deviates slightly from *f* (*y*) when the non-linearity is strong (*n* = 10, Fig S2A and B), which results from the fact that the time-average of *f* (*y_t_*) is not equivalent to 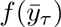, an effect that becomes pronounced when *n* is large.

We also examined a bell-shaped fitness landscape, confirming that 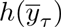 recovered *f* (*y*) to good precision (see Materials and Methods and Fig S3 in detail). These results show that our lineage-based analysis allows us to probe fitness and selection strength of heterogeneous cellular phenotypes from realistic sample sizes of single-cell lineage trees.

We emphasize that it is relatively rare to find cases in which population growth is driven by the instantaneous value of a single measurable phenotype, such as *y_t_* above. That is, one should generally not equate the fitness landscape extracted for a single phenotype, 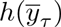, with the overall physiological fitness landscape of cells, a much more complex, multi-dimensional quantity. Instead, *h*(*y*_*τ*_) constitutes the effective fitness landscape for the phenotype of interest, 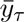, and despite the underlying complexity of cellular physiology, it remains well defined and experimentally measurable.

### Single-cell time-lapse experiment

Next, we apply the analytical framework to analyze single-cell time-lapse data of *E. coli* cells that express an antibiotic resistance gene *sm*^*R*^ [31] and a fluorescent reporter *venus-yfp* [32]. We constructed and used two strains in the experiments: F3/pTN001, in which *venus* and *sm*^*R*^ are transcribed together under the control of a common promoter PLlacO-1 [33] on a low copy plasmid pTN001 (pSC101 ori), but translated separately (Fig 5A); and F3NW, in which the fusion protein, Venus-SmR is expressed from the *intC* locus on the chromosome under the control of PLlacO-1 promoter (Fig 5B). The Sm^R^ protein confers resistance to a ribosome-targeting antibiotic drug, streptomycin, by direct inactivation [34, 35]. We conducted fluorescent time-lapse measurements of cells proliferating on agarose pads that contain either no drug (−Sm) or a sub-inhibitory concentration of streptomycin (+Sm) (200 µg/ml for F3/pTN001; and 100 µg/ml for F3NW) (Fig 5C). The minimum inhibitory concentrations (MIC) of streptomycin for F3/pTN001 and F3NW were 1000 µg/ml and 250

**Fig 5.**
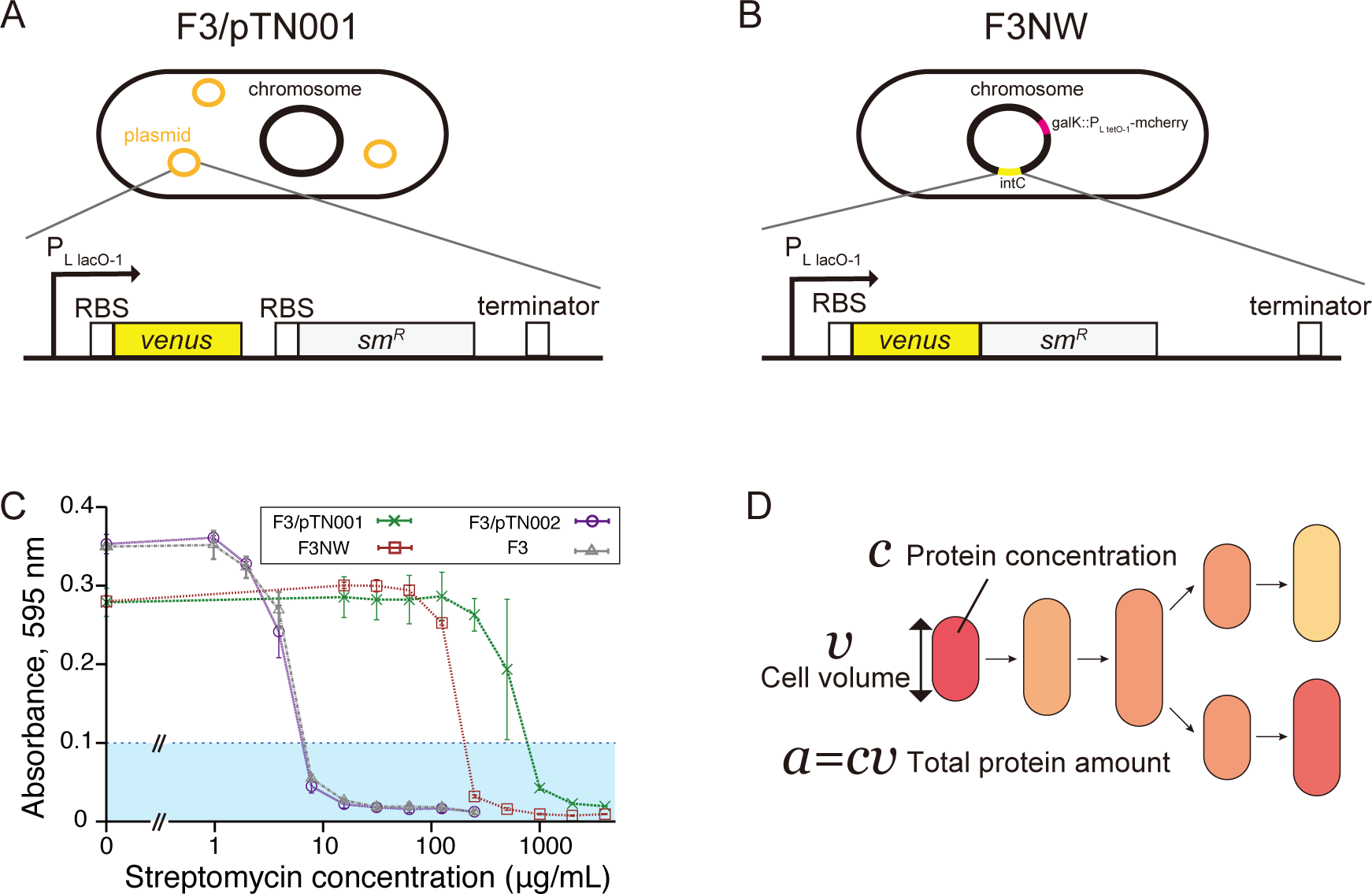
*E. coli* strains used in single-cell time-lapse measurements. **A.** F3/pTN001 strain. This strain expresses a fluorescent protein Venus-YFP and streptomycin Venus-Sm^R^ under the control of the PLlacO-1 promoter from *intC* locus on the chromosome. To facilitate the image analysis, we additionally integrated an mcherry-rfp gene that is expressed under the control of the PLtetO-1 promoter [33] from *galK* locus on the chromosome. **C.** MICs of streptomycin for F3, F3/pTN001, F3/pTN002, and F3NW. Absorbance at 595 nm of cell cultures of each strain at different concentrations of streptomycin was measured after 20-hour incubation with shaking at 37° C. pTN002 is a negative control plasmid in which the *sm*^R^ gene was removed from pTN001. The average of three replicates are plotted with the standard deviation for each condition of streptomycin concentration. We determined MICs by the minimum concentration above which the absorbance of cell culture remains below 0.1 (cyan region): 8 µg/mL for F3 (gray) and F3/pTN002 (purple), 250 µg/mL for F3NW (brown), and 1000 µg/mL for F3/pTN001 (green). **D.** Quantities obtained from time-lapse images. We extracted the time-series of cell volume *v*, protein concentration (mean fluorescence intensity per cell area) *c*, and total protein amount (sum of fluorescence intensity of the pixels within a cell) *a* = *cv* together with cell lineage trees, and calculated 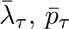, and 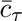 for each cell lineage according to the definitions in Eq. 15–17.

µg/ml, respectively, which were significantly higher than the MIC of the parental strain F3 (8 µg/ml) (Fig 5C). Thus, Sm^R^ protein is functional in the constructed strains. We extracted the information of lineage trees along with time-series of cell size *v*(*t*) and of fluorescence intensity *c*(*t*) (Fig 5D). Since *c*(*t*) is a proxy for protein concentration in a cell, *c*(*t*)*v*(*t*) can be regarded as the quantity that scales with the total amount of protein in a cell. Based on these quantities, we analyzed three different time-averaged phenotypes along a single-cell lineage: elongation rate 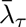, protein production rate 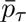, and protein concentration 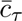, which are defined as

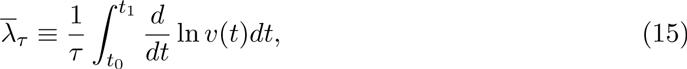

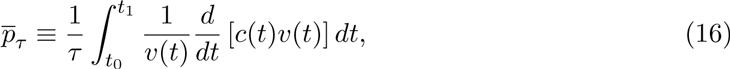

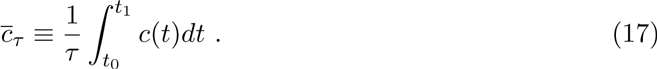

We calculated these phenotypic quantities for all the lineages spanning from *t*_0_ to *t*_1_, andobtained the chronological probability distribution *P*^cl^(·), fitness landscape *h*(·), and selection strength *S*[·] of these phenotypes.

### Fitness landscapes and selection strength of F3/pTN001

We first analyzed the growth of F3/pTN001. Population growth kinetics revealed that the growth rate difference between −Sm and +Sm conditions was small and became noticeable only after *t* = 200 min (Fig 6A). Therefore, we focused on the time window between *t*_0_ = 200 min and *t*_1_ = 400 min (see Fig S4 for the results when *t*_0_ = 0 min and *t*_1_ = 200 min). The population growth rates during this period were 0.45±0.01 h^*−*1^ for −Sm and 0.39±0.01 h^*−*1^ for +Sm, respectively (*p <* 0.05) (Fig 6B). Consistently, the mean of lineage fitness in the chronological perspective 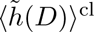 in +Sm condition was 0.35±0.01 h^*−*1^, which is smaller than that in −Sm condition, 0.41±0.01 h^*−*1^ (*p <* 0.05) (Fig 6C). Despite the decrease in the mean lineage fitness, we did not detect the difference in intra-population lineage heterogeneity measured by maximum selection strength *S*[*D*] (*p* = 0.5) (Fig 6D).

**Fig 6.**
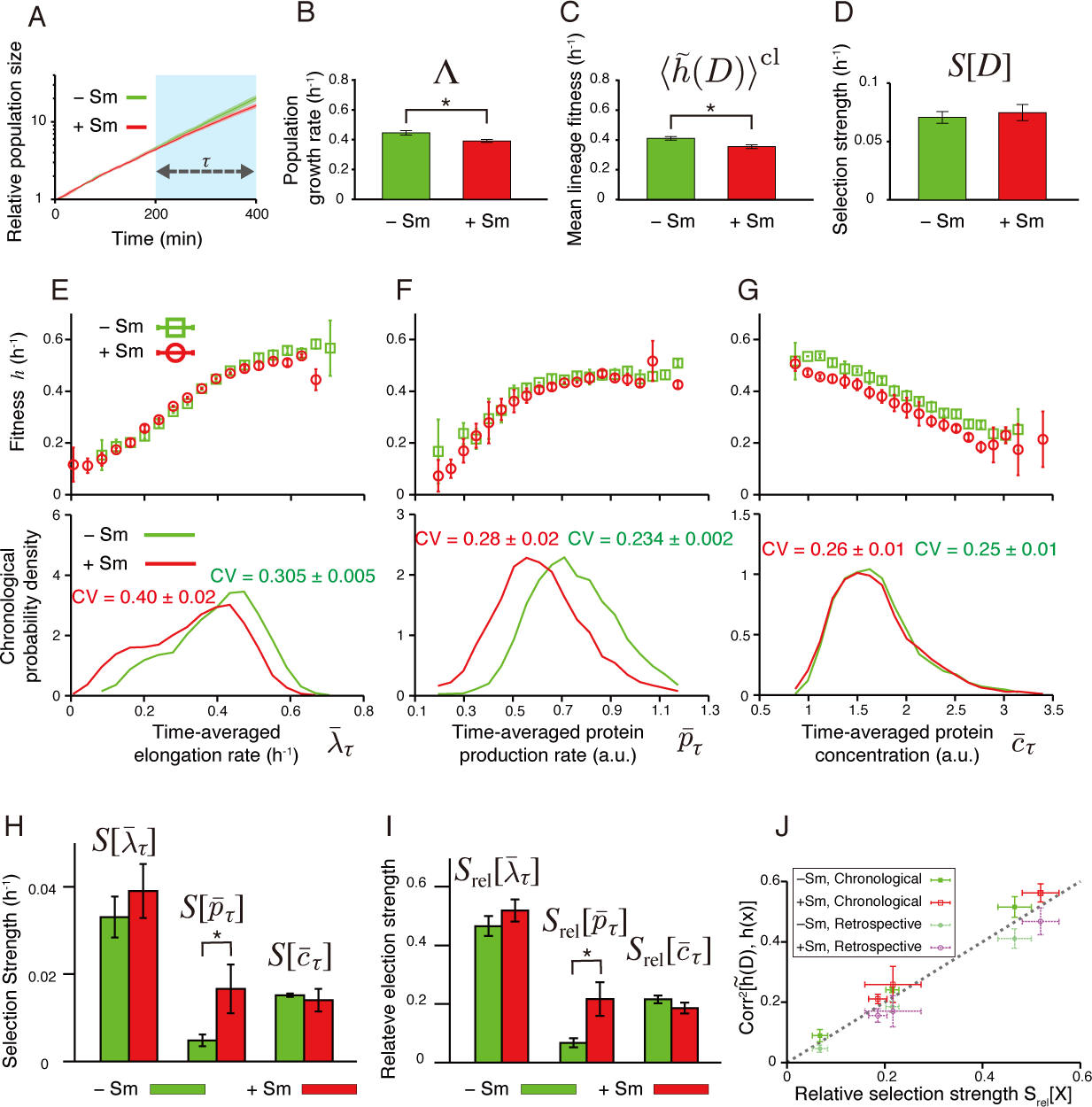
Fitness landscapes and selection strength measured for *E. coli* F3/pTN001. **A.** Population growth curves. Green curve is for −Sm condition, and red for +Sm condition (the color correspondence is the same for all subsequent panels). Relative population size on the y-axis is the number of cells at each time point normalized by the number of cells at *t* = 0 min. The error bars in all panels are the standard deviations of three independent experiments. Growth rate difference became apparent only after *t* = 200 min. Hence, we set *t*_0_ = 200 min and *t*_1_ = 400 min in the following analyses. The results with *t*_0_ = 0 min and *t*_1_ = 200 min are shown in Fig S4. The numbers of cells at times *t*_0_ and *t*_1_, which specify the number of cell lineages used in the analysis, are given in Table S1. **B-D.** Comparison of population growth rate Λ (B), chronological mean lineage fitness 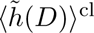 (C), and selection strength for division count *S*[*D*] (D), between −Sm and +Sm conditions. *p*-values by *t*-test are 0.013, 0.010, and 0.529, respectively (*n* = 3). **E-G.** Fitness landscapes *h*(*x*) (upper panels) and chronological distributions *P*^cl^(*x*) (lower panels) for elongation rate 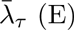, protein production rate 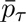 (F), and protein concentration 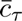 (G). The fitness landscapes for elongation rate and protein production rate were barely distinguishable between −Sm and +Sm conditions, whereas that for protein concentration shows a slight downshift in +Sm condition. In contrast, shift of chronological distributions was observed for elongation rate and protein production rate, but not for protein concentration. **H.** Selection strengths. We compared selection strengths 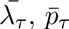 and 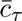between −Sm and +Sm conditions, finding a statistically significant difference only for 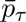 (*p <* 0.05). The *p*-values are 0.34 for 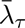, 0.044 for 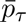, and 0.58 for 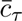, respectively (*n* = 3). **I** Relative selection strengths. Again, the difference is statistically significant only for *<*inline-formula>. The *p*-values are 0.21 for 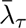, 0.024 for 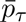, and 0.14 for 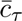, respectively (*n* = 3). **J.** Relationship between relative selection strength and squared correlation coefficient between 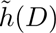 and *h*(*x*), where 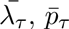, or 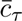. The correlation coefficients were evaluated by both chronological and retrospective probabilities.

The three lineage phenotypes had distinct characteristics in their response to the drug (Fig 6E–I). The fitness landscapes of elongation rate were nearly identical between −Sm and +Sm conditions, and increased approximately linearly with 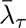 (Fig 6E and Fig S5A, B). This agrees with the natural assumption that fast elongation should lead to proportionately high division rate. The chronological distribution 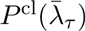 shifted to the left in +Sm condition (Fig 6E), which is also consistent with the fact that 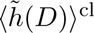 is slightly lower in +Sm condition. Nevertheless, we did not detect the difference in selection strength 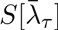 (Fig 6H). These results confirm that 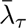 behaves coherently with *D* under these conditions.

The fitness landscape of protein production rate were likewise nearly identical between +Sm and −Sm conditions (Fig 6F and Fig S5C, D). The landscape is a more saturating function rather than linear with the kink around 0.5 a.u.. The fact that 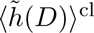 is an increasing function even in the absence of the drug is presumably because overall cellular metabolism couples to all production rates and cells growing faster generally have higher production rates in most genes. The chronological distribution 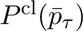 shifted significantly toward the left in +Sm condition. Interestingly, we detected an increased selection strength 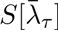 in +Sm condition (1.7 × 10^*−*2^ h^*−*1^) compared with that in −Sm condition (0.5 × 10^*−*2^h ^*−*1^, *p <* 0.05) (Fig 6H). The relative selection strength 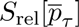 was also significantly different (Fig 6I). Because 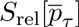 is a measure of correlation between 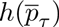 and lineage fitness 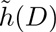, resistance-conferring protein Sm^R^ under the control of the PLlacO-1 promoter from a low copy plasmid pTN001 (pSC101 ori). Ribosomal binding sites are present in front of the start codons of both structural genes, thus proteins are translated separately. We analyzed the data assuming that production rate and protein amount of Sm^R^ are strongly correlated with those of Venus-YFP. **B.** F3NW strain. This strain expresses a fusion protein this result indicates that the heterogeneity in Sm^R^ production rate becomes more strongly correlated with fitness in +Sm condition than in −Sm condition. This change in the selection strength largely comes from the shift of the chronological distribution 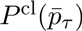: A large portion of the probability distribution resides in the plateau region of the fitness landscape in –Sm condition, whereas its shift in +Sm condition causes a significant overlap with the linear region, resulting in a larger fitness heterogeneity in the phenotypic space of 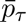.

The fitness landscapes of protein concentration decrease linearly with 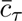 in both +Sm and −Sm conditions; protein expression levels and fitness are thus anti-correlated (Fig 6G and S5E, F). Surprisingly, we did not detect any advantages of high expression level even in the presence of the drug (Fig 6G). The chronological distribution 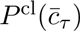 and selection strength 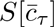 were nearly identical between the two conditions (Fig 6G and 6H). This indicates that, unlike production rate 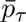 the strength of correlation between Sm^R^ expression level and fitness is unchanged even if the drug is added. The results therefore suggest that the protein production rate of Sm^R^ is a more responsive phenotype to drug than protein expression level in this strain. The response characteristics of selection strength are unchanged even if the relative selection strengths were compared between the two conditions (Fig 6I).

Applying fitness decomposition in Eq. 13 to the experimental data revealed that the changes of mean fitness in retrospective perspective due to the environmental change from −Sm to +Sm (Δₓ*h*(*x*)〉^rs^) mostly came from the changes in Δₓ*h*(*x*)〉^cl^, not from the changes in selection strengths Δ*S*[*x*], for all the phenotypes (Table 1). Therefore, the contribution of Δ*S*[*x*] to Δₓ*h*(*x*)〉^rs^ were marginal at least in the environmental difference used in this study. We found that the relative selection strengths of 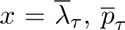, and 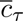 were approximately equal to the squared correlation coefficients between 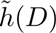 and *h*(*x*) evaluated by both chronological and retrospective probabilities (Fig 6J). This validates the simple interpretation that 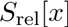 represents the correlation between 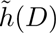 and *h*(*x*), though the small differences of the squared correlation coefficients between the chronological and retrospective probabilities suggest the contribution of higher-order cumulants (*Supporting Information*).

**Table 1.** Contributions of individual cells’ response Δ〈*h*(*x*)〉^**cl**^ **and change of selection strength** Δ*S*[*x*] **to fitness gain** 〈*h*(*x*)〉^rs^ for the environmental change from −Sm to + Sm.

| | Division $D$ | Elongation $\bar{\lambda}_{\tau}$ | Protein production $\bar{p}_{\tau}$ | Protein concentration $\bar{c}_{\tau}$ |
| --- | --- | --- | --- | --- |
| $\Delta\langle h(x) \rangle^{\text{rs}}$ | $-0.05 \pm 0.02$ | $-0.05 \pm 0.02$ | $-0.05 \pm 0.02$ | $-0.06 \pm 0.02$ |
| $\Delta\langle h(x) \rangle^{\text{cl}}$ | $-0.06 \pm 0.02$ | $-0.06 \pm 0.02$ | $-0.06 \pm 0.02$ | $-0.06 \pm 0.02$ |
| $\Delta S[x]$ | $0.005 \pm 0.009$ | $0.007 \pm 0.008$ | $0.012 \pm 0.006$ | $0.000 \pm 0.004$ |

### Fitness landscapes and selection strength of F3NW

We next examined how the difference in the expression scheme between F3/pTN001 and F3NW affected the phenotype distributions, fitness landscapes, and selection strength (Fig 7). For F3NW, we focused on the time window between *t*_0_ = 100 min and *t*_1_ = 300 min, where the difference in population growth rate was significant (0.52± 0.02 h^*−*1^ in −Sm condition, and 0.48 ± 0.01 h^*−*1^ in +Sm condition) (Fig 7A and 7B). We did not detect statistically significant differences in 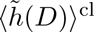 (0.49± 0.03 h^*−*1^ in −Sm, and 0.45 ± 0.01 h^*−*1^ in +Sm, Fig 7C) and in *S*[*D*] (0.06 ± 0.01 h^*−*1^ in −Sm, and 0.057 ± 0.004 h^*−*1^ in +Sm, Fig 7D).

**Fig 7.**
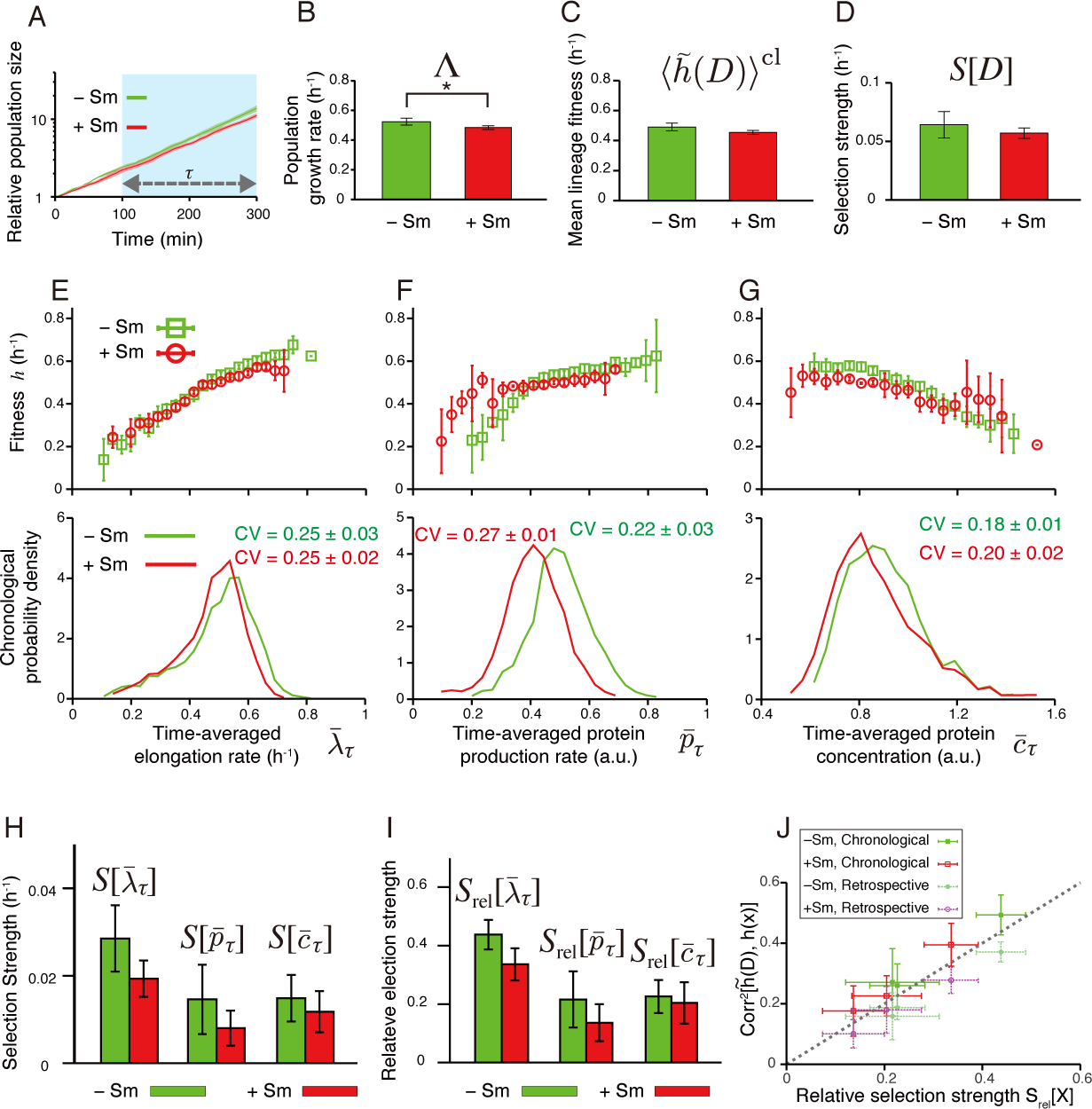
Fitness landscapes and selection strength measured for *E. coli* F3NW. **A.** Population growth curves. Green curve is for −Sm condition, and red for +Sm condition (the color correspondence is the same for all the following panels). Relative population size on the y-axis is the number of cells at each time point normalized by the number of cells at *t* = 0 min. The error bars in all panels are the standard deviations of three independent experiments. Growth rate difference became apparent only after *t* = 100 min. Hence, we set *t*_0_ = 100 min and *t*_1_ = 300 min in the following analyses. The numbers of cells at time *t*_0_ and *t*_1_ are shown in Table S2. **B-D.** Comparison of population growth rate Λ (B), chronological mean lineage fitness 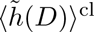, and selection strength for division count *S*[*D*] (D), between −Sm and +Sm conditions. *p*-values by *t*-test are 0.036, 0.077, and 0.34, respectively (*n* = 4). **E-G.** Fitness landscapes *h*(*x*) (upper panels) and chronological distributions *P*^cl^(*x*) (lower panels) for elongation rate 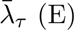 (E), protein production rate 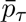, and protein concentration 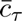. **H.** Selection strengths. We compared selection strengths 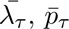, and 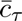 between −Sm and +Sm conditions, finding no statistically significant differences for all the phenotypes. The *p*-values are 0.12 for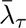, 0.25 for 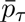, and 0.48 for 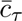, respectively (*n* = 4). **I.** Relative selection strengths. Again, no statistically significant differences were found. The *p*-values are 0.057 for 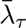, 0.27 for 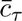, and 0.69 for *<*inline-formula> respectively (*n* = 4). **J.** Relationship between relative selection strength and squared correlation coefficient between 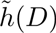 and *h*(*x*), where 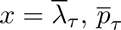, or 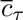. The correlation coefficients were evaluated by both chronological and retrospective probabilities.

Comparing the fitness landscapes between the two strains revealed that the overall shapes of the fitness landscapes were unchanged by the difference of the expression schemes (Fig 7E–7G and Fig S6A-F): For elongation rate of F3NW strain, fitness landscapes increased with 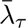 almost linearly; for production rate, fitness increases monotonically with 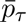 with a kink; and for protein concentration, fitness decreases monotonically with 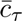. One important difference is that the fitness landscape of protein production rate 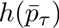 in +Sm condition shifted significantly toward the left along with the distribution (Fig 7F). Consequently, the main part of the distribution of 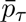 remained in the range where the fitness is fairly uniform (Fig 7F), and the selection strength 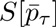 of F3NW did not increase in +Sm condition (Fig 7H and 7I). The selection strength of 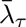 and 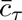 of F3NW was also unchanged between –Sm and +Sm conditions (Fig 7H and 7I) as seen in F3/pTN001 (Fig 6H and 6I).

The measured selection strength of the three phenotypes was close to the squared correlation coefficients between *h*(*x*) and *h*˜(*D*) (Fig 7J), which again validates the interpretation that 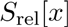 is a measure of correlation between phenotype *x* and fitness.

Interestingly, we found that the chronological distributions of the three phenotypes of F3NW were all narrower than those of F3/pTN001 (Fig 7E–7G, and S7). This indicates that expressing Sm^R^ and Venus from the plasmid induced additional heterogeneity in all the phenotypes including elongation rate. Such non-trivial effects of different gene expression schemes can be also probed by this method quantitatively. We note that the ranges where we can assess the fitness landscapes became narrower in F3NW than in F3/pTN001 simply because of the lower levels of phenotypic heterogeneity of this strain.

## Discussion

Phenotypes of individual cells are intrinsically heterogeneous, and phenotypic heterogeneity is ubiquitously seen across taxa from microbes to mammalian cells. different phenotypic states among genetically identical cells can be selected within a population when they are correlated with fitness. Therefore, unraveling the unique phenotypic characteristics that allowed a certain set of cell lineages to outperform in a population is important for understanding the biological roles of the phenotypic heterogeneity of interest.

We have presented a method to quantify fitness differences and selection strength for heterogeneous phenotypic states of individual cells within a population. Our framework shares a basic idea with the method for measuring selection strength developed in evolutionary biology in that we evaluate phenotype-dependent fitness [36–38]. The key novelty of our approach is that we consider individual lineages or histories as the basic units of proliferation. An important advantage of this history-based formulation of fitness landscapes and selection strength is that it is applicable even to cellular phenotypes that fluctuate in time, such as gene expression levels in single cells. Indeed, we demonstrated by simulation that the pre-assigned fitness landscape could be recovered from the single-cell lineage trees and the associated dynamics of cellular phenotypic states despite the stochastic transitions of internal, cellular states. Though a number of single cell studies have suggested the functional roles of phenotypic fluctuation in a genetically uniform cell population [5, 6], our framework provides the first procedure for the rigorous quantification of the fitness values of such fluctuating cellular states. In this framework, we can use any statistical quantities that are measurable on cell lineages as the ‘phenotype’. Although we exclusively evaluated the time-averages of cellular phenotypes along cell lineages in the analysis, other statistical quantities such as variance and coefficient of variation can also be evaluated as lineage phenotypes, which might reveal e.g. the fitness value of “noisiness” of gene expression level. Conversely, the flexibility imposes a technical challenge to select a suitable quantity that correctly reports cellular functions. We emphasize that the fitness landscapes and selection strengths quantified in this study report only correlation between the lineage phenotypes and cell division, not causality. To address causality, one must carefully choose appropriate lineage phenotypes that take detailed time-series of phenotypic states into account.

One of the key features of our analysis is that we measure fitness by cell divisions, and not by cell size growth such as elongation rate, which is widely used as a proxy for fitness in single-cell analysis on bacterial growth. There are two important reasons for this: (1) population growth is ultimately driven by cell divisions, not by cell elongation; and (2) selection strength *S*[*D*] imposes the fundamental upper limit on the strength of selection for any phenotype. We are interested in determining which single-cell variables are under the strongest selection, which can be assessed by our measure of relative selection strength, *S*_rel_[*x*] = *S*[*x*]*/S*[*D*]. Thus, evaluating the fitness and selection through the correlation with cell divisions has a fundamental importance for studying selection in a population.

We applied our method to the clonal proliferation processes of *E. coli*, and quantified the fitness landscape and the selection strength for different phenotypes with and without an antibiotic drug. First, we found that the elongation rate was the phenotype with largest relative selection strength, across conditions and strains, with values of 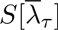 that ranged from 30% to 50% of the maximum possible value. This indicates that elongation rate behaves like a trait that is under strong phenotypic selection within clonal populations of *E. coli*. As mentioned previously, this analysis on its own cannot determine the causal relations, i.e. whether elongation rate is directly under selection, or indirectly by correlation with another trait. All other things being equal, however, cells that elongate faster are likely to divide sooner, and their lineages will thus be amplified with respect to cells that elongate slower and divide later, yielding a simple mechanism for the selection we detect.

Second, we made an interesting observation concerning the selection strength for the time-averaged protein concentration of Sm^R^, which was indistinguishable between the two environments with and without the drug, whereas that for time-averaged protein production rate increased significantly by drug exposure in F3/pTN001. This result indicates that, at least for this particular strain and experimental condition, the production rate is a more responsive phenotype that increases its correlation with fitness in +Sm condition. This does not mean that Sm^R^ protein concentration is less important for the fitness in +Sm condition. The correlation between phenotype and fitness in each condition is represented by *S*_rel_[*x*] itself, not by the change in *S*_rel_[*x*]. 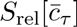 of F3/pTN001 remains at a high level both in −Sm and +Sm conditions (Fig 6I), thus its heterogeneity is significantly correlated with the lineage fitness. It is, however, surprising that the heterogeneity in Sm^R^ protein concentration is not correlated with fitness in the +Sm condition any more than that in the −Sm condition, considering the known functional role of Sm^R^ protein in inactivating the drug. It would be important for the future studies to examine the fitness landscapes and selection strength for broader sets of drug conditions and resistance proteins.

Our method characterized the similarities and differences of phenotype distributions, fitness landscapes, and selection strength between the closely related *E. coli* strains (F3/pTN001 and F3NW) (Fig 7). Interestingly, the results revealed that the phenotypes of F3NW were all less heterogeneous than those of F3/pTN001 (Fig S7). This suggests that even a small difference of expression scheme could affect the heterogeneity levels of a large set of phenotypes. Even if one knows that different strains have different levels of phenotypic heterogeneity, the consequences for fitness are usually difficult to evaluate rigorously. Our method extracts such information from experimentally obtainable lineage trees and the measured transition of phenotypes along cellular lineages.

We emphasize that our method evaluates net results, i.e. the fitness landscapes and selection strength for each phenotype whose heterogeneity can be caused by many possible noise sources. The fitness landscapes and selection strength of F3/pTN001 and F3NW are themselves valid for describing the properties of the measured phenotypes in each strain, and comparing the results among the strains would provide the contributions of different noise sources such as plasmid copy number variations. The same is true for cases where multiple cell types coexist in a population due to cellular differentiation or bet-hedging; even when the differences between these cell types are not easily apparent, our method can evaluate the overall fitness landscapes and selection strength of the phenotypes of interest. When one can clearly identify differences between cell types using markers, such a parameter can be directly incorporated into the analysis as a phenotype, and the contribution of coexisting cell types to the overall population growth can then be unraveled.

Recently, several groups have demonstrated that the heterogeneity of division intervals in clonal cellular populations increases population growth rate [15, 39]. Cerulus, *et al*. showed that the levels of variability and epigenetic inheritance of division intervals are changeable to a large extent depending on the environments and the genetic backgrounds in *Saccharomyces cerevisiae*. In general, larger variations and stronger epigenetic inheritance of division times cause stronger selection in the population, and our method allows us to quantify the contribution of these factors to population growth by the selection strength *S*[*D*]. Since *S*[*D*] is directly measurable using lineage trees, and has a clear meaning as the upper bound of selection strength for any phenotype, the statistics of division counts have a fundamental importance in our lineage analysis framework.

Conventional genetic perturbation methods such as gene knock-out, overexpression, and gene suppression only associate a population-level gene expression state with population fitness; they are unable to report whether different expression states of single cells in the same population are correlated to their fitness. Our new analytical framework, however, allows us to reveal the impact of different expression levels and dynamics on cellular fitness without modifying population-level expression states, and might open up a new field in genetics that connects different expression states to cellular fitness without applying the genetic perturbation.

The application of this method is not restricted to the analysis of clonal proliferation in unicellular organisms. An important application would be in the analysis of embryogenesis and stem cell differentiation of multicellular organisms, in which cellular reproduction rates diversify among the branches of lineage trees as the differentiation process goes forward [40]. Recently, large-scale cell lineage trees along with detailed quantitative information on cellular phenotypes (gene expression, cell position, movement, etc.) have been available [19,20,24,25]. Quantifying fitness and selection strength for different phenotypes at the single-cell level in differentiation processes might reveal key phenotypic steps and events leading to cell fate diversification. Additionally, fruitful applications may be found in the analysis of evolutionary lineages in viral populations, such as influenza [41] and HIV [42], where lineage trees have been obtained using temporal sequencing data. Quantifying the strength of selection on viral traits, such as antigenic determinants, and inferring their fitness landscape is an important challenge in the field [43–45] which the method presented here could address. The application of this new lineage analysis tool to broader biological contexts may unravel the roles of phenotypic heterogeneity in diverse cellular and evolutionary phenomena.

## Materials and Methods

### Simulation

We simulated clonal cell proliferation processes using a custom C program. We determined phenotypic state *y_t_*_+Δ*t*_ by randomly sampling the value of ln *y_t_*_+Δ*t*_ from the normal distribution with mean *µ* + *e*^*−γ*^^Δ*t*^ (ln *y_t_* − *µ*) and variance *σ*^2^ (1 − *e*^*−*2*γ*Δ*t*^) assuming that the transition of ln *y_t_* follows the Ornstein-Uhlenbeck process. We set Δ*t* = 5 min, *µ* = −0.5 ln(1.09), *σ*^2^ = ln(1.09), and *γ* = (−0.6 ln *r_g_*) h^*−*1^ with *r_g_* = 0.8. In this setting, *y_t_* follows the log-normal distribution with mean 1.0 and standard deviation 0.3 in the stationary state without selection (i.e. Hill coefficient *n* = 0). We assumed that cells divide with the probability of *f* (*y_t_*)Δ*t* where 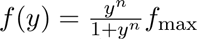 with f_max_= 1.2 h*−*1 at each time point, and the initial states of two daughter cells (*y_t_*_+Δ*t*_) were determined independently of each other from the last state (*y_t_*) of their mother cell. Without selection, the division rate is *f*_0_ = *f*_max_/2 = 0.6 h^*−*1^ and thereby the mean interdivision time along a lineage is 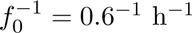. Without selection, since the normalized autocorrelation function of ln *y_t_* at stationary sate is *φ*(*τ*) = *e*^*−γτ*^, *r_g_* = *e*^*−γ/f*^0 is the autocorrelation of ln *y_t_* after a single generation. Fitness landscapes of *y*_*τ*_ with faster fluctuation conditions (*r_g_* = 0.5 and 0.2) were shown in Fig S2. We also ran the simulations with another type of instantaneous reproduction rate 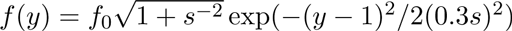 with *f*_0_ = 0.6 h and with *s* = 0.5, 1, 2 (Fig S3). We produced a dataset that contains 100 lineage trees (i.e. *N* (*t*_0_) = 100 cells) with the length of *τ* = 250 min in each condition, which is comparable to the data size of the real experiments (Table S1). For each condition, we repeated the simulation 10 times, and the average and standard deviation were shown in Fig 4 and in Fig S2.

### Cell strains and culture conditions

We used F3, F3/pTN001, F3/pTN002, and F3NW *E. coli* strains in the experiments. F3 is a W3110 derivative strain in which three genes (*fliC, fimA*, and *flu*) are deleted. pTN001 and pTN002 are low copy plasmids constructed from pMW118 (Nippon Gene, Co., LTD). We constructed pTN001 by introducing the P_LlacO-1_ promoter [33], *venus* gene [32], *sm*^R^ gene, t1t2 *rrnB* terminator, and frt-franked kanamycin resistance cassette [46] into the multi-cloning site of pMW118. We also placed ribosome-binding sites in front of both *venus* and *sm*^R^ genes; these two genes are transcribed together, but translated separately. pTN002 is a control plasmid that lacks the *sm*^R^ gene from pTN001. F3NW expresses the fusion protein of Venus-Sm^R^ from the *intC* locus of the chromosome under the control of the P_LlacO-1_ promoter. We also introduced *mcherry-rfp* into the *galK* locus on the chromosome under the P_LtetO-1_ promoter [33] to facilitate the microscopic observation and image analysis. See Supplemental

Information and Table S3 for the details on how we constructed these plasmids and strains. We cultured the cells in M9 minimal medium (M9 minimal salt (Difco) + 2 mM MgSO_4_ (Wako) + 0.1 mM CaCl_2_ (Wako) + 0.2% glucose (Wako)) at 37°C. 0.1 mM Isopropyl *β*-D-1 thiogalactopyranoside (IPTG) (Wako) was added to the cultures of F3/pTN001, F3/pTN002, and F3NW to induce the expression of the genes under the control of the P_LlacO-1_ promoter. For single-cell time-lapse experiments, we solidified M9 medium with 1.5% (w/v) agarose (Gene Pure Agarose, BM Bio). We adjusted the IPTG concentration in M9 agarose to 0.1 mM by adding ×1,000 concentrated IPTG solution to the melted M9 agarose before solidification. Approximately 5 mm (W)×8 mm (D)×5 mm (H) piece of M9 agarose gel was mounted onto cell suspension on a glass-bottom dish (IWAKI). For +Sm condition, we added 200 µg/mL streptomycin when solidifying M9 agarose gel.

### Determination of MIC

Overnight cultures of the four *E. coli* strains in M9 medium at 37°C from glycerol stock were diluted ×100 into 2-ml fresh M9 medium and cultured for three hours at 37°C. 100 µl exponential phase culture was mixed with 100 µl fresh M9 medium containing streptomycin in a 96-well plate. We prepared 10 different conditions of streptomycin concentration for each strain with the concentration increased in two-fold stepwise. The optical density of the cell cultures after mix was ca. 0.05 at 600 nm. The cell cultures in a 96-well plate were incubated by shaking at 37°C for 20 hours. We determined the MICs of F3 and F3/pTN001 with a microtiter plate (FilterMax F5, Molecular Devices) by absorbance at 595 nm.

### Time-lapse microscopy

To prepare a sample for time-lapse microscopy, we first cultured the cells from glycerol stock in M9 medium at 37°C by shaking overnight. Next, we diluted the overnight culture ×100 in 2 ml fresh M9 medium, and cultured it for another three hours at 37_*°*_C by shaking. We adjusted the OD_600_ of the culture to 0.05, and 1 µl of the diluted culture was spread on a 35 mm (*×*) glass-bottom dish (IWAKI) by placing M9 agarose pad onto the cell suspension. To avoid drying the M9 agarose pad, water droplets (total 200 µl) were placed around the internal edge of the dish. The dish was sealed by parafilm to minimize water evaporation. Fluorescent time-lapse images were acquired every 5 minutes with Nikon Ti-E microscope equipped with a thermostat chamber (TIZHB, Tokai Hit), 100x oil immersion objective (Plan Apo *λ*, N.A. 1.45, Nikon), cooled CCD camera (ORCA-R2, Hamamatsu Photonics), and LED excitation light source (DC2100, Thorlabs). The temperature around the dish was maintained at 37°C. The microscope was controlled by micromanager (https://micro-manager.org/).

### Analysis

Time-lapse images were analyzed with a custom macro of Image J (http://imagej.nih.gov/ij/). This macro produces the results file, which contains the information of mean fluorescence intensity, cell size (area), and geneaological position of individual cells. We analyzed the results file with a custom C program.

To evaluate fitness landscapes and selection strengths both in the simulation and the experiments, we determined the bin width based on the interquartile range of each phenotypic state (Fig S8-10). The details are explained in *Supporting Information*.

## Acknowledgments

This work was supported by JSPS KAKENHI Grant Number 25711008, 15KT0075, 15H05746 (YW); NIH Grant Number R01-GM-097356 (EK); and Platform for Dynamic Approaches to Living System from MEXT and AMED, Japan (YW). TN was supported by Grant-in-Aid for JSPS Fellows (14J01376). We thank Reiko Okura and Sayo Akiyoshi for technical assistance, Atsushi Miyawaki for providing Venus/pCS2 plasmid, Hironori Niki for providing pKP2375 plasmid, Ippei Inoue for the technical advice on *E. coli* strain and plasmid construction, and members of Wakamoto lab for in-depth discussions.

